# MEGaNorm: Normative Modeling of MEG Brain Oscillations Across the Human Lifespan

**DOI:** 10.1101/2025.06.23.660997

**Authors:** Mohammad Zamanzadeh, Ymke Verduyn, Augustijn de Boer, Tomas Ros, Thomas Wolfers, Richard Dinga, Marie Šafář Postma, Andre F. Marquand, Marijn van Wingerden, Seyed Mostafa Kia

## Abstract

Normative modeling provides a principled framework for quantifying individual deviations from typical brain development and is increasingly used to study heterogeneity in neuropsychiatric conditions. While widely applied to structural phenotypes, functional normative models remain underdeveloped. Here, we introduce MEGaNorm, the first normative modeling framework for charting lifespan trajectories of resting-state magnetoencephalography (MEG) brain oscillations. Using a large, multi-site dataset comprising 1,846 individuals aged 6–88 and spanning three MEG systems, we model relative oscillatory power in canonical frequency bands using hierarchical Bayesian regression, accounting for age, sex, and site effects. To support interpretation at multiple scales, we introduce Neuro-Oscillo Charts, visual tools that summarize normative trajectories at the population level and quantify individual-level deviations, enabling personalized assessment of functional brain dynamics. Applying this framework to a Parkinson’s disease cohort (n = 160), we show that normative deviation scores reveal disease-related abnormalities and uncover a continuum of patients in theta–beta deviation space. This work provides the first lifespan-encompassing normative reference for MEG oscillations, enabling population-level characterization and individualized benchmarking. All models and tools are openly available and designed for federated, continual adaptation as new data become available, offering a scalable resource for precision neuropsychiatry.

## 1 Introduction

Normative modeling [1, 2] has emerged as a transformative approach in neuroimaging, providing a robust framework to characterize individual deviations from typical brain structure and function by establishing population-level reference distributions. Unlike traditional case-control studies that rely on group-averaged comparisons, which often mask biologically meaningful variability, normative models enable subject-specific quantification of deviations. This approach captures the continuous nature of brain variation across demographic and clinical populations [3]. It has been particularly impactful in structural magnetic resonance imaging (sMRI) studies, where it has been used to map deviations in patients with neuropsychiatric disorders using various imaging-derived phenotypes (IDPs), such as cortical thickness [4], subcortical volumes [5], and white matter microstructure [6, 7]. For example, in schizophrenia [8, 9] and autism spectrum disorder [10], normative models have uncovered distinct neuroanatomical alterations and identified subgroups of patients with atypical developmental trajectories that remain undetected using conventional group-based analyses [11]. Similarly, in Alzheimer’s disease, normative approaches have helped disentangle age-related atrophy from disease-specific pathological changes [12, 13]. By providing individualized deviation profiles, normative modeling represents a crucial step toward precision medicine [14, 15], offering a data-driven framework for understanding disease heterogeneity and informing personalized diagnostic and therapeutic strategies.

While normative models of sMRI have shown considerable promise in advancing precision neuropsychiatry, extending this framework to functional neuroimaging modalities is essential for deepening our understanding of complex brain disorders and diseases. Many neuropsychiatric conditions are defined by abnormal functional brain dynamics, which can be studied using techniques such as functional MRI (fMRI), electroencephalography (EEG), and magnetoencephalography (MEG) [16–18]. Normative modeling has already been applied to fMRI data to establish normative maps of functional brain dynamics [19–21]. However, normative models for EEG and MEG data are still underdeveloped, despite their high temporal resolution and ability to capture neuronal dynamics [22–24], reflecting key functional processes including sensory integration, attention, and memory [25, 26].

Understanding how brain oscillations evolve across the lifespan has long been a central objective in neuroscience. Early efforts in this area focused on EEG and introduced foundational approaches to characterize normative brain activity [27–29]. While these studies pioneered the concept of normative baselines for electrophysiological signals, they relied on relatively simple parametric models and were often constrained by small or demographically narrow samples. More recently, advances in data acquisition and processing have enabled more detailed lifespan analyses using both EEG [30–32] and MEG [33–36]. For example, Rempe et al. [36] quantified age-related changes in relative and absolute power across the lifespan in resting-state MEG (rs-MEG), while Tröndle et al. [37] examined alpha-band power development in adolescents using data from the Human Brain Network. However, these recent studies have focused on modeling average age-related trends and have not accounted for population-level variability through centile-based modeling. In addition, they rely on a single dataset, sometimes with restricted age ranges, which constrains generalizability across scanner types and the full human lifespan.

In this study, we introduce the MEGaNorm framework, an end-to-end rs-MEG processing pipeline for 1) deriving lifespan normative ranges of functional imaging-derived phenotypes (f-IDPs) on large, multiscanner MEG datasets, and 2) mapping individuals’ f-IDPs to their deviations from the norm of the population. Using this framework, we present the first normative model of rs-MEG oscillations, derived from the relative power spectrum of theta, alpha, beta, and gamma frequency bands after removing the aperiodic component of the signal. By isolating periodic activity, our normative models capture agerelated spectral power changes of neural oscillations, independent of broadband power shifts in absolute power [38]. The models are constructed using large-scale rs-MEG data from multiple scanners and hardware vendors. To account for this inter-site variability, we employ hierarchical Bayesian regression (HBR) [4, 39] that enables the simultaneous estimation of non-Gaussian and heteroscedastic age-specific centiles of variation in brain oscillations while modeling site-specific effects.

We further show how the derived normative models can be used to systematically characterize the changes in lifespan trajectories of resting-state brain oscillations at both population and individual levels. At the population level, we estimate growth charts of relative spectral powers with different trajectories for male and female populations across the lifespan. Using these growth charts, we derive Population-level Neuro-Oscillo Charts (P-NOCs), which show how the relative distribution of brain oscillations in theta, alpha, beta, and gamma frequency bands changes across the human lifespan. At the individual level, we introduce Individual-level Neuro-Oscillo Charts (I-NOCs), which map individual spectral power measurements onto population norms, conditioned on age, sex (following the SAGER guidelines [40], we use the term “sex” limited to “female” and “male”), and scanning device. I-NOCs enable a fine-grained assessment of individual deviations, offering a personalized interpretation of brain dynamics concerning the normative range of brain oscillations in a reference population. Finally, to demonstrate the translational utility of our framework, we benchmark our model on a cohort of patients with Parkinson’s disease, leveraging normative deviation scores to quantify oscillatory abnormalities within the patient population. We show how normative models can help conceptualize a neuropsychiatric disorder as a continuum of abnormalities in f-IDPs, marking a crucial step toward precision neuropsychiatry.

## 2 Results

We developed *MEGaNorm*, a normative modeling framework to chart lifespan trajectories of MEG brain oscillations and to identify individual deviations from these normative ranges. We used rs-MEG recordings from 1,846 healthy individuals aged 6–88 and a clinical cohort of 160 individuals with Parkinson’s disease aged 43–88. These data were pooled from six independent datasets acquired using three major MEG systems (see Section 4.1), offering broad coverage of the human lifespan and diverse acquisition environments. This dataset was used to derive reference models of f-IDPs, defined as sensor-level relative power in canonical frequency bands.

To generate these normative models, MEGaNorm integrates a sequence of processing and modeling steps (Figure 1). MEG recordings were first preprocessed to remove artifacts and segmented into fixed-length epochs. Power spectral densities (PSDs) were then computed and decomposed into periodic and aperiodic components to isolate oscillatory activity. Relative power in the theta, alpha, beta, and gamma bands was extracted from the periodic activity and averaged across sensors, yielding 4 device-agnostic f-IDPs. Hierarchical Bayesian regression (HBR)[4, 41, 39] was applied to model lifespan trajectories, accommodating non-Gaussian distributions, heteroscedasticity, and site- and sex-specific effects. This framework enabled the estimation of age-conditioned centiles for each frequency band, providing the foundation for visualizing normative developmental trends and assessing individual deviations. Full methodological details are provided in the Methods Section 4.

**Figure 1:**
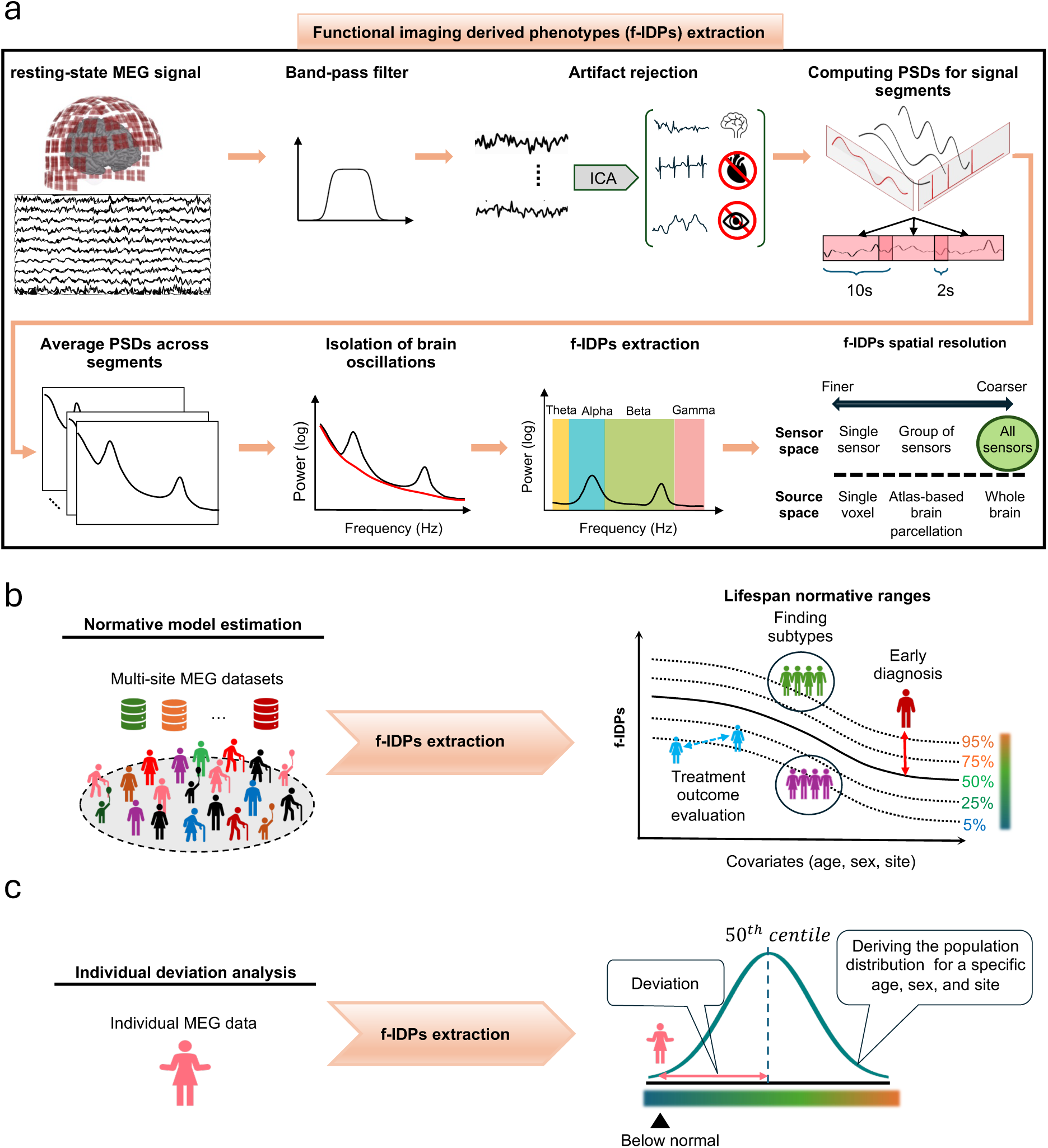
Overview of the MEGaNorm pipeline. MEGaNorm is an end-to-end framework for deriving normative models of resting-state MEG (rs-MEG) oscillations and quantifying individual deviations from population-level norms. The pipeline includes the extraction of functional imaging-derived phenotypes (f-IDPs), modeling normative trajectories across the lifespan, and characterizing individual-level deviations. **(a)** Signal preprocessing and f-IDP extraction: Raw rs-MEG signals undergo bandpass filtering and artifact removal using independent component analysis (ICA). Data are segmented into fixed-length epochs, and power spectral densities (PSDs) are computed and averaged across segments. Periodic activity is isolated by separating aperiodic components, and relative power is extracted in canonical frequency bands. These band-specific features are then averaged across sensors to obtain device-agnostic f-IDPs. **(b)** Normative model estimation: f-IDPs extracted from a large, multi-site cohort are used to estimate lifespan trajectories using hierarchical Bayesian regression. These models yield age-specific normative centiles for each frequency band, accounting for site and sex effects. The resulting normative ranges can support clinical applications such as early diagnosis, treatment monitoring, and subtyping of neuropsychiatric disorders. **(c)** Individual deviation analysis: For a new individual, f-IDPs are computed and compared against normative models conditioned on age, sex, and recording site. This enables the quantification and visualization of individualized deviations from normative trajectories.

### 2.1 Model evaluation: modeling non-gaussianity, heteroscedasticity, and non-linearity results in more accurate centiles

We evaluated the quality of the fitted normative models using both quantitative and qualitative out-of-sample diagnostics. We performed an ablation study comparing our proposed non-linear, heteroscedastic, and non-Gaussian HBR model (see section 4.4 for details) to a simplified baseline. The proposed model used a B-spline basis for age, a Sinh-Arcsinh (SHASH) likelihood [42, 39], and modeled variance as a function of age. In contrast, the baseline model employed a linear age term, a Gaussian likelihood, and assumed homoscedastic variance. This comparison aimed to determine whether incorporating flexible distributional assumptions improves the fidelity of normative centile estimates.

Both models were evaluated using repeated 50%–50% train–test splits of healthy participants, stratified by site (see Section 4.5 for details). Model training and evaluation were repeated 10 times using different random seeds for train–test splits to assess the stability and generalizability of performance. A range of evaluation metrics was used to assess both the location and shape of the fitted distributions. To quantify the accuracy of the fitted curve at the median (50^th^ percentile), we computed the standardized mean squared error (SMSE), which evaluates how well the model predicts the central tendency of the data. To evaluate the distributional properties of the derived *z*-scores as a measure of model calibration, we calculated skewness, excess kurtosis, and the Shapiro–Wilk test statistic (*W*)[43]. To directly assess the accuracy of centile estimation, we used the mean absolute centile error (MACE), introduced in this study for evaluating the calibration of estimated centiles (see Section 4.5 for more information). Results are summarized in Figure 2. As a complementary qualitative assessment, Q–Q plots were used to visually evaluate the alignment between empirical and theoretical Gaussian quantiles.

**Figure 2:**
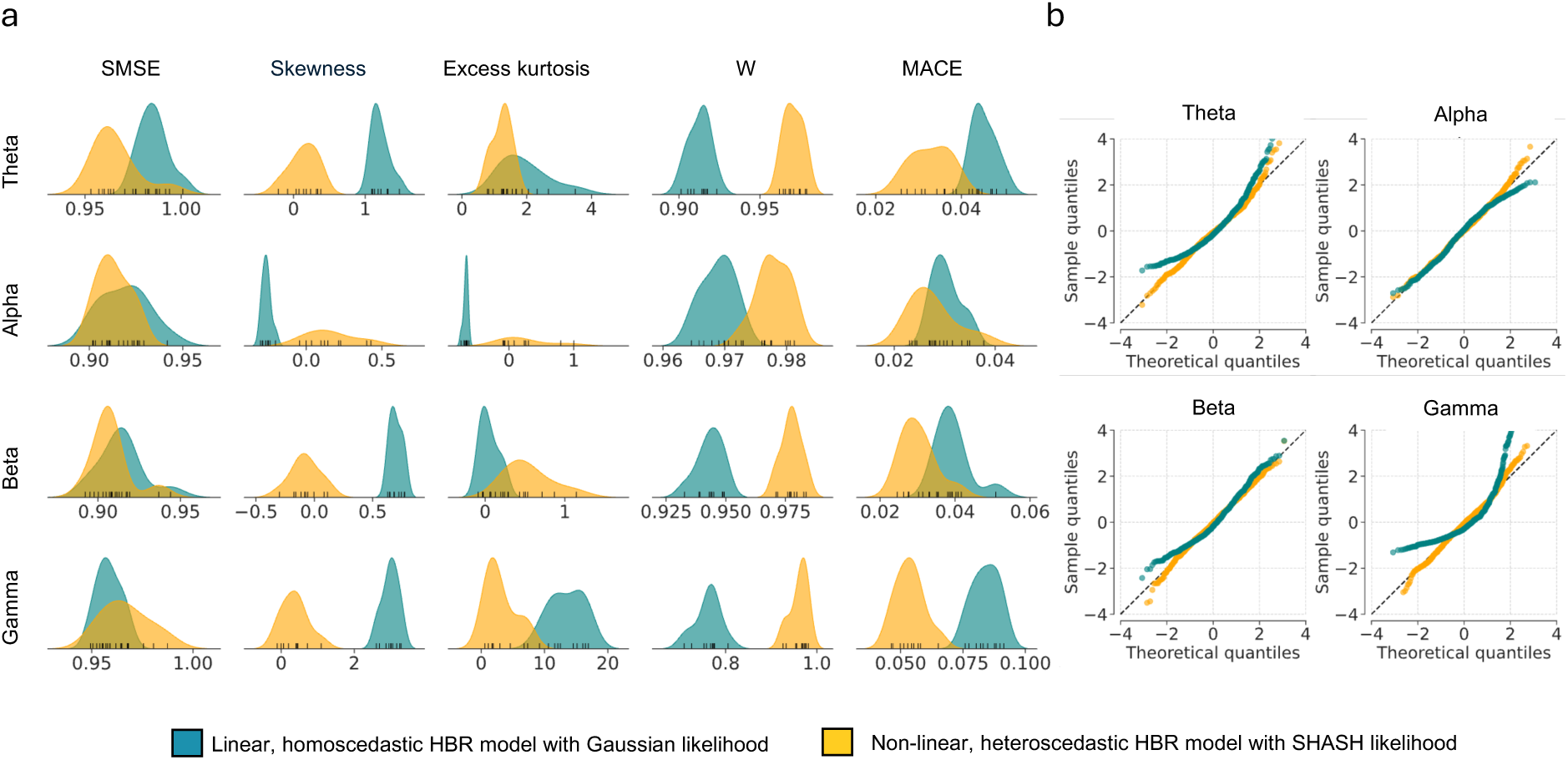
Model evaluation and comparison. **(a)** Summary of out-of-sample model diagnostics for four frequency-specific f-IDPs: standardized mean squared error (SMSE), skewness, excess kurtosis (kurtosis minus 3), Shapiro–Wilk test statistic *W*, and mean absolute centile error (MACE). Lower SMSE values reflect a more accurate estimation of the true central tendency with the baseline at 1. Skewness and excess kurtosis values closer to zero, along with W values closer to 1, reflect greater normality of the z-score distributions. Lower MACE values indicate better centile calibration. Here, the MACE was evaluated at the 1^st^, 5^th^, 25^th^, 50^th^, 75^th^, 95^th^, and 99^th^ centiles. Across all f-IDPs, the proposed non-linear, heteroscedastic, and non-Gaussian HBR model outperformed the baseline in modeling the distribution of response variables. (b) Q–Q plots of z-scores from the held-out test set. The black dashed 1:1 line represents the theoretical quantile–quantile correspondence under a standard Gaussian distribution. The proposed model shows closer adherence to Gaussianity compared to the baseline.

Although both models performed similarly in predicting the central tendency, quantified by SMSE, with significant improvement observed only in the theta band (see Table 1), the proposed HBR model provided substantially better estimates of the full distributional shape. Across all frequency bands, *z*-score distributions derived from the proposed model more closely approximated a standard Gaussian distribution, as reflected in *W* statistics closer to 1.0 and skewness and excess kurtosis values nearer to zero. While mild leptokurtosis remained, the distributions exhibited improved symmetry and tail behavior. These improvements were visually corroborated by Q–Q plots (Figure 2b), which showed tighter alignment between empirical and theoretical quantiles. Additionally, the proposed model achieved significantly lower MACE values across all frequency bands (Wilcoxon rank-sum test *p <* 0.05), indicating superior centile calibration. Together, these findings suggest the proposed model more accurately captures the shape of the underlying distributions, an essential feature for deriving well-calibrated normative centiles. This supports the inclusion of non-Gaussianity, heteroscedasticity, and non-linearity in modeling the f-IDPs.

**Table 1:**
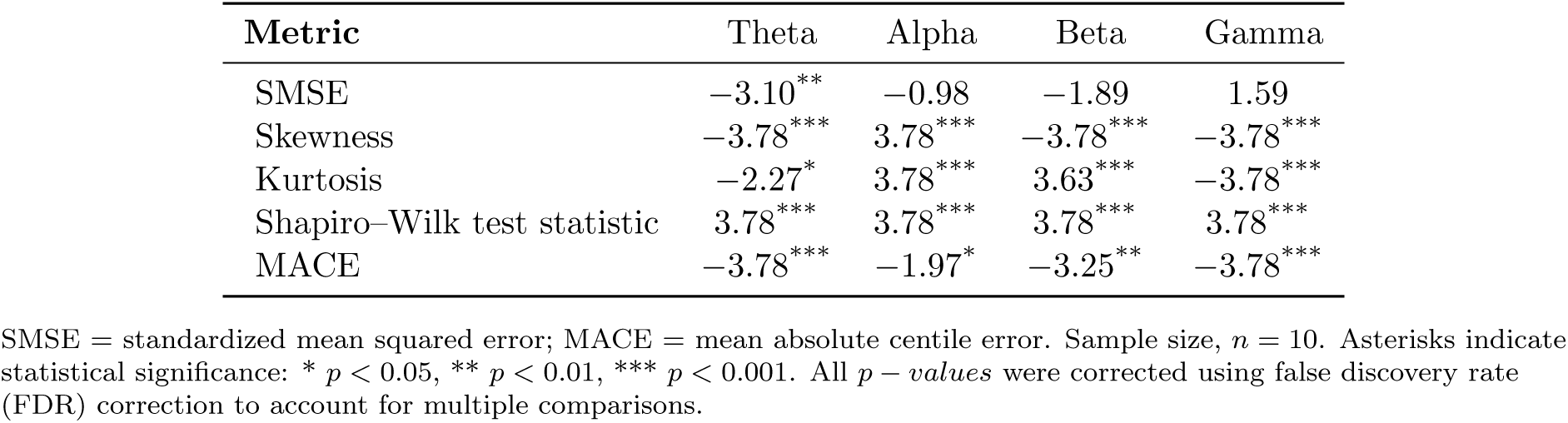
*W* statistics of Wilcoxon rank-sum test comparing the proposed (non-linear, heteroscedastic, and non-Gaussian) with the baseline HBR model (linear, homoscedastic, and Gaussian).

### 2.2 Population- and individual-level Neuro-Oscillo Charts

Building on the accuracy of the proposed HBR model, we used the full dataset of healthy individuals to derive growth charts for four oscillatory f-IDPs. These charts were generated by sampling from the posterior predictive distribution across ages 6 to 80, separately for males and females (Figure 3a). The resulting centile trajectories reveal substantial inter-individual variability and exhibit complex age-related dynamics, including heteroscedasticity and non-linear trends.

**Figure 3:**
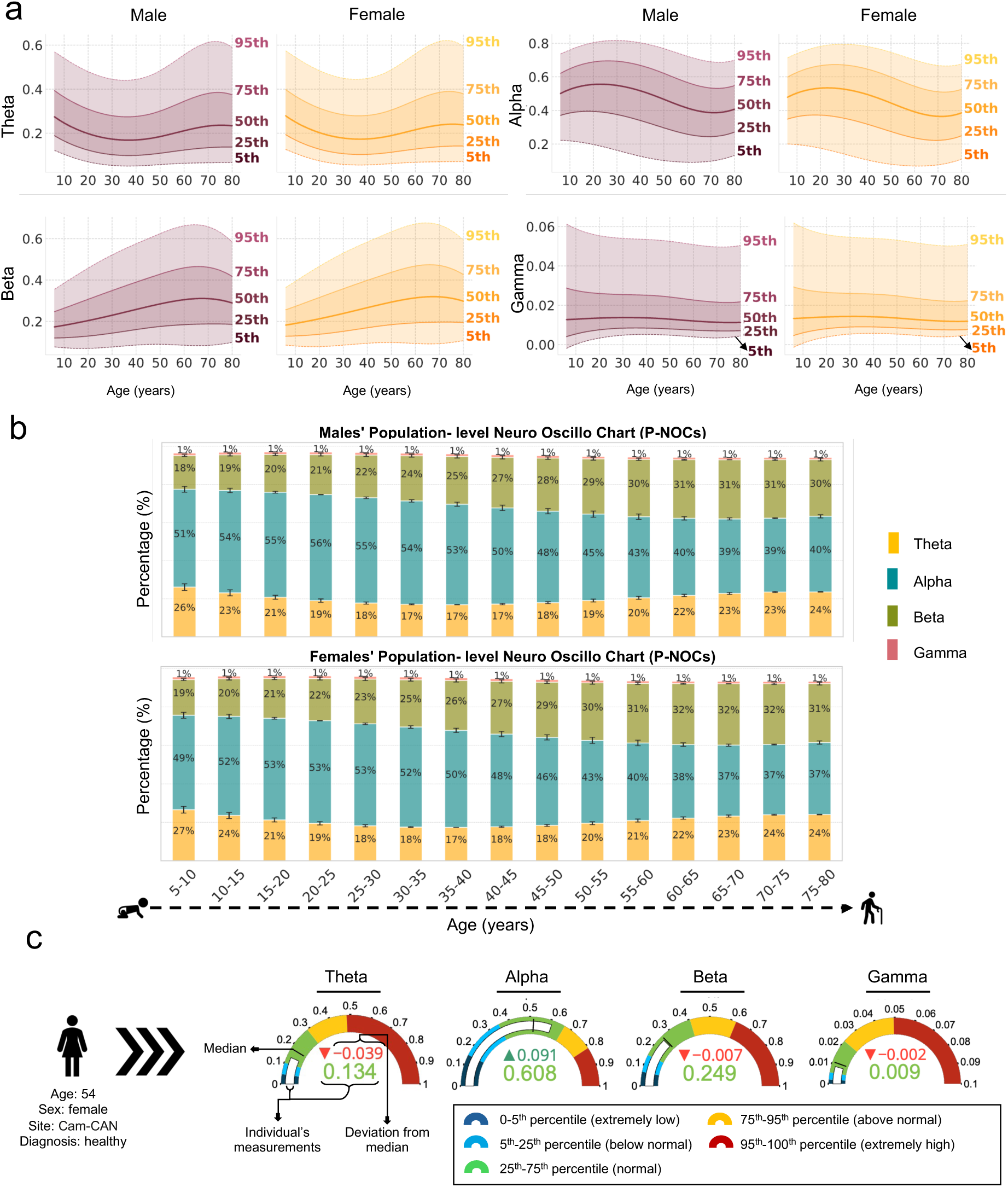
From growth charts to population- and individual-level Neuro-Oscillo Charts. **(a)** Lifespan growth charts for each frequency band (theta, alpha, beta, and gamma), stratified by sex. Centiles were derived by sampling from the posterior predictive distribution of the estimated HBR model. For visualization purposes, centiles were averaged across datasets. **(b)** Population-level Neuro-Oscillo Charts (P-NOCs), which depict the distribution of relative power across frequency bands at the 50^th^ centile, separately for males and females. P-NOCs summarize how much the spectral power of each frequency band contributes to the total oscillatory power and how it evolves with aging, enabling intuitive comparisons of population-level trends across sex and lifespan. **(c)** Individual-level Neuro-Oscillo Charts (I-NOCs), which visualize a participant’s spectral power relative to age-, sex-, and site-matched normative distributions derived from the growth charts. The white bar represents the participant’s measurement, which is also shown by numerical values below the gauge plot (color-coded by the respective range). The upper value below the gauge plots displays the participant’s measurement deviation from the 50^th^ percentile (shown by black line). This example uses data from a clinically undiagnosed 54-year-old female participant from the Cam-CAN dataset.

To summarize normative developmental patterns at the population level, we introduce Population-level Neuro-Oscillo Charts (P-NOCs), which visualize age-related changes in the relative power of frequency bands for 5-year intervals (ages 5 to 80) at the median (50^th^) centile. P-NOCs illustrate the relative contribution of each frequency band’s 50*^th^* centile to the total power spectra. Figure 3b shows P-NOCs for male and female populations. These charts reveal key lifespan trends: theta power follows a U-shaped trajectory with a minimum in midlife (ages 30–40), alpha power follows an inverted U-shaped trend peaking in early adulthood (ages 20–30), beta power increases steadily with age, and slow gamma remains relatively constant, albeit small in magnitude.

To enable individualized interpretation and enhance the clinical utility of the model, we further introduce Individual-level Neuro-Oscillo Charts (I-NOCs), which allow single-subject benchmarking against normative trajectories. I-NOCs visualize an individual’s f-IDP values in relation to the reference normative range, conditioned on the participant’s demographic information. Figure 3c illustrates I-NOCs for a healthy participant, conditioned on the participant’s age, sex, and recording site (a 54-year-old female participant from the Cam-CAN dataset), whose values fall within the normal range across all frequency bands. For example theta’s I-NOC shows that theta power accounts for 13.4% of the total oscillatory power, which is 3.9% below the population median (50th percentile), yet remains within the normal range (i.e., between 25^th^ to 75^th^ percentiles). I-NOCs offer an intuitive and scalable tool for personalized f-IDP profiling, providing a foundation for real-time downstream applications in anomaly detection and patient stratification.

### 2.3 Deviations from normative ranges are informative in Parkinson’s disease

We evaluated the clinical utility of the derived normative models in an anomaly detection framework by assessing whether large deviations from normative ranges could identify individuals with Parkinson’s disease. This was done using models trained exclusively on data from healthy individuals, without any exposure to patients’ data [4]. Specifically, we used the OMEGA dataset, selecting 221 undiagnosed participants from the test partition as controls and 160 Parkinson’s disease patients from the same dataset as the clinical cohort.

For each participant, deviation scores (z-scores) were computed and converted into abnormal probability indices using the method introduced in Kia et al. [4] (see also Methods 4.6 for details). We then evaluated the discriminative power of each frequency-specific f-IDP by calculating the area under the receiver operating characteristic curve (AUC). Statistical significance (*p <* 0.05) was assessed using permutation testing (1000 repetitions), with false discovery rate (FDR) correction for multiple comparisons [44]. This procedure was repeated across 10 different random train-test splits of the healthy cohort, while the same Parkinson’s disease cohort was used in each iteration.

Figure 4a summarizes AUC results across ten runs. Among the four f-IDPs, theta z-scores showed the strongest discriminative performance (mean AUC = 0.62, SD = 0.01), significantly outperforming chance in all iterations. Beta z-scores also demonstrated significant predictive power (mean AUC = 0.59, SD = 0.01) across all runs. In contrast, gamma z-scores (mean AUC = 0.53, SD = 0.02) were significant in only 2 out of the 10 runs, and alpha z-scores (mean AUC = 0.45, SD = 0.008) failed to reach significance in any run. These findings suggest that deviations in theta and beta band power are informative markers of disease-related functional abnormalities.

**Figure 4:**
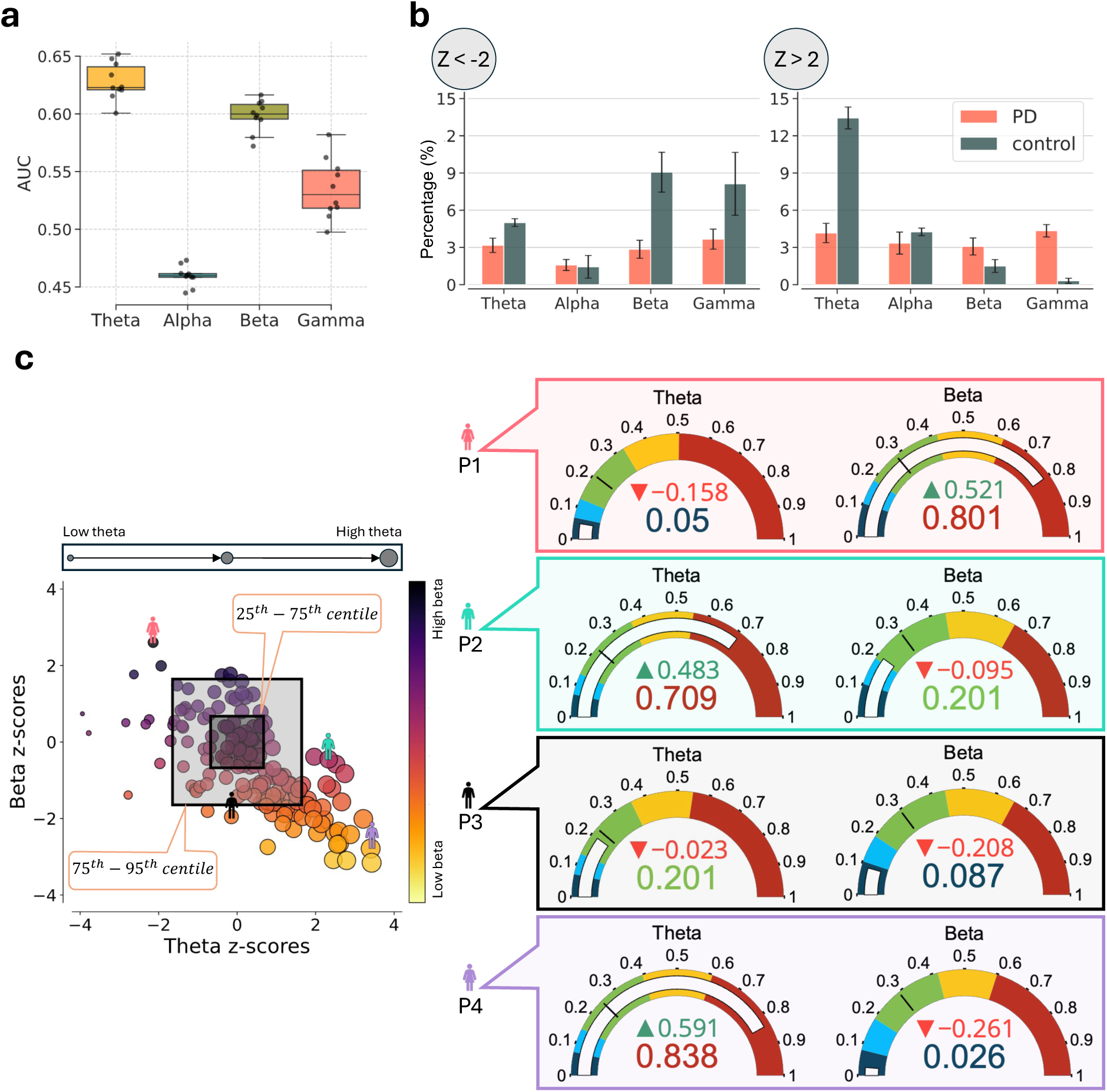
Individual-level analysis in the clinical population with Parkinson’s disease. **(a)** Area under the ROC curves (AUCs) for anomaly detection across ten independent train–test splits. Z-score deviations were used to compute abnormal probability indices for each frequency band. **(b)**Percentage of participants exhibiting positive (*Z >* 2) and negative (*Z < −*2) extreme deviations across frequency bands, shown separately for healthy controls and patients. Bars represent averages over ten iterations; error bars indicate 95% confidence interval. **(c)** Scatter plot of patients in theta–beta deviation space. Marker size encodes theta z-scores (x-axis), and color encodes beta z-scores (y-axis). Selected individuals (P1–P4) illustrate distinct regions of this spectrum and are accompanied by their corresponding I-NOCs.

### 2.4 Decoding heterogeneity in Parkinson’s disease via theta-beta deviation profiles

To further characterize the directionality and heterogeneity of functional abnormalities in Parkinson’s disease, we examined the distribution of extreme deviations (*|Z| >* 2) in f-IDPs among patients. As shown in Figure 4b, 13.43% of patients exhibited positive extreme deviations (*Z >* 2) in the theta band, while 9.06% showed negative extreme deviations (*Z < −*2) in the beta band. Additionally, 8.12% of patients displayed negative deviations in gamma power. Deviations in the alpha band, by contrast, were infrequent and not substantial.

To explore whether these deviations reveal underlying subtypes, we visualized the joint distribution of patients in theta–beta deviation space, the two most informative frequency bands identified in the anomaly detection analysis (see Supplementary results for theta-gamma and beta-gamma deviation spaces). As illustrated in Figure 4c, this space reveals a continuum of oscillation profiles ranging from high-theta/low-beta to low-theta/high-beta, capturing the heterogeneity of disease-related abnormalities. Notably, a subset of patients falls within the interquartile range for both bands, indicating the absence of marked deviations.

To illustrate this variability more concretely, we present I-NOCs for four representative patients with Parkinson’s disease (P1–P4) located at different points along the theta–beta deviation spectrum. These I-NOCs reveal that patients diagnosed with the same condition can exhibit markedly different neurophysiological profiles, falling in different quantile ranges of the norm. Patients P1 and P4 are positioned toward opposite extremes of the spectrum: P1 shows extremely low relative theta power (5.0%) and extremely high beta power (80.1%), whereas P4 exhibits the opposite pattern, with extremely high theta (83.8%) and extremely low beta (2.6%). In contrast, patients P2 and P3 display large deviations in only one of the frequency bands. P3’s beta power (8.7%) is extremely low, while its theta deviation falls within the normal range. Conversely, P2 has relatively normal beta power (20.1%, within the interquartile range), but extremely elevated theta. These examples highlight the utility of normative modeling in capturing individual variability across Parkinson’s disease patients.

## 3 Discussion

In this study, we addressed the long-standing challenge of delineating normative developmental trajectories of brain oscillations across the human lifespan [45, 27, 28, 30, 33, 34, 36, 46–48]. To this end, we introduced *MEGaNorm*, a normative modeling framework for estimating age-related centiles of f-IDPs based on large-scale, multi-site rs-MEG data covering ages 6 to 88. The derived models offer a transdiagnostic solution by providing normative reference distributions not constrained by diagnostic categories, thereby supporting applications across a wide range of neuropsychiatric conditions. To demonstrate clinical relevance, we applied MEGaNorm to a Parkinson’s disease cohort and showed that deviation scores captured individual-specific functional abnormalities that would be obscured in conventional case-control analyses. These results are consistent with the dimensional approach that emphasizes characterizing mental disorders along continuous axes of neurobiological variation. In the following, we outline key methodological advances introduced by MEGaNorm that extend the current state of the art and offer substantial value for both basic and translational neuroscience.

To enhance the specificity of the derived oscillatory f-IDPs, we isolated periodic brain oscillations from the aperiodic component of the MEG signal. Electrophysiological recordings inherently contain both oscillatory (periodic) activity and a broadband aperiodic background, which includes neural and non-neural contributions such as instrumental noise and physiological artifacts [37, 49, 38]. Aperiodic features have been shown to vary systematically with age [50–52], arousal [53], and individual factors like educational background [54], and may be confounded by cardiac artifacts that also exhibit age-dependent variation [52]. Failing to account for these influences can obscure true oscillatory dynamics. By separating these components before feature extraction, MEGaNorm yields a more accurate and interpretable quantification of spectral power, thereby supporting a more reliable characterization of normative brain function across the lifespan.

To ensure generalizability and robust estimation of normative trajectories, we constructed our models using a large, multi-site dataset comprising recordings from multiple MEG hardware systems. This design choice allowed us to better capture population-level variability and improve the applicability of the derived centiles across diverse acquisition settings. The estimation of reliable centiles in normative modeling, particularly when using flexible methods such as HBR, critically depends on large sample sizes [5, 55]. To address this need, we aggregated six independent rs-MEG datasets spanning a wide age range and three different MEG manufacturers. While this multi-site strategy introduces valuable diversity, it also presents challenges due to scanner and protocol-specific differences, environmental noise, and sample heterogeneity [56]. We mitigated these effects by modeling the site as a batch variable in HBR using partial pooling [4], enabling the model to share information across sites while retaining sensitivity to systematic inter-site differences (see Supplementary results). This principled handling of site effects enhances the robustness and transferability of the resulting normative models.

We adopted a flexible normative modeling framework based on HBR to accommodate non-Gaussian distributions, heteroscedasticity, and nonlinear age-related trends. This methodological flexibility was critical for producing accurate, data-driven growth charts that capture both the central tendency and dispersion of f-IDPs across the lifespan. Earlier studies have reported inter-individual variability in the healthy aging of brain oscillations using descriptive tools such as box plots [33], confidence intervals [57], or by noting wide deviations around mean trajectories [58, 35]. Such variability is likely shaped by diverse biological and environmental influences, including genetic factors, neurodevelopmental differences, and contextual exposures [59, 60]. Capturing this variability requires models that not only estimate the mean developmental trajectory but also account for the distributional shape and its evolution with age. Empirical findings have demonstrated that the variance and skewness of oscillatory power can vary substantially with age [33, 61], and that transformations are often needed to approximate Gaussianity in EEG datasets [62]. Moreover, studies have shown that nonlinear trends, such as quadratic patterns, are better suited for modeling changes in alpha and gamma power across the lifespan [35, 36]. To address these challenges, we employed a semi-parametric modeling strategy using B-spline basis functions, which enables flexible estimation of nonlinear trajectories without imposing rigid parametric assumptions [63]. Our findings confirmed that the resulting growth charts accurately reflect non-Gaussian, heteroscedastic variation in f-IDPs.

The MEGaNorm framework is inherently extensible and designed to accommodate calibration on new, local, or private datasets [4]. As additional data become available, posterior distributions from pre-trained models can be leveraged as informative priors, enabling model refinement without requiring re-estimation from scratch. This feature is particularly valuable for sites with limited sample sizes or restricted age ranges, as it enables partial pooling across datasets while preserving sensitivity to site-specific effects. Notably, prior work has shown that even small calibration cohorts, as few as 25 participants, are sufficient to adapt HBR-based models to new populations [64]. Crucially, this adaptation does not necessitate centralized data sharing: the hierarchical structure of HBR supports federated learning, allowing models to be updated locally while maintaining data privacy and regulatory compliance. To facilitate broader adoption, the data processing pipeline and derived models are openly shared, respectively via the MEGaNorm package (v0.1.0) [65] and PCNPortal [66], supporting reproducibility and enabling continual improvement as the community contributes new data.

We also introduced two complementary visualization tools, P-NOCs and I-NOCs, that enhance the interpretability of normative modeling outputs at both the population and individual levels. P-NOCs depict how the relative contributions of canonical frequency bands proportionally evolve with age. These charts serve as compact summaries of developmental and aging trajectories, facilitating population-level comparisons across demographic groups. In contrast, I-NOCs provide subject-specific benchmarking by situating an individual’s spectral power profile within the normative distribution, conditioned on individuals’ demographic information such as age, sex, and recording site. I-NOCs enable personalized assessments that are especially useful in clinical applications, including early diagnosis and monitoring of treatment effects.

To demonstrate the clinical utility of MEGaNorm, we applied the framework to a cohort of individuals with Parkinson’s disease, a condition known to involve widespread disruptions in neural oscillatory dynamics [67–70]. A well-documented feature of Parkinson’s disease is neural slowing, typically characterized by elevated power in lower-frequency bands (e.g., theta and alpha) and reduced power in higher-frequency bands (e.g., beta and gamma) [71, 72]. However, these patterns are inconsistently observed across studies, with variability attributed to differences in patient subgroups[71], disease stages [73, 74], and methodological factors. In particular, some studies report no significant differences in alpha or beta power between patients and controls [75, 76], and the effects of dopamine replacement therapy on oscillatory power remain equivocal [74, 77]. These discrepancies likely reflect the clinical and biological heterogeneity of Parkinson’s disease [78, 79], as well as the limitations of conventional case-control analyses that rely on group-averaged contrasts, which can obscure meaningful inter-individual variation [11, 80]. Normative modeling offers a principled alternative by estimating individual-level deviations from a population-based reference, thus enabling the characterization of functional abnormalities along a continuous spectrum rather than dichotomous case-control categories [3, 1].

Using the normative modeling approach within an anomaly detection framework [4], we identified significant deviations in theta and beta frequency bands among individuals with Parkinson’s disease, with average AUCs of 0.62 and 0.59, respectively. Importantly, these models were trained in an unsupervised manner on healthy controls only and were not tailored to disease-specific features. This unsupervised and transdiagnostic nature underscores the generalizability of the framework to other neuropsychiatric populations. Despite not being disease-optimized, the models successfully captured meaningful deviations related to Parkinson’s disease pathology.

To further investigate the structure of abnormalities in theta-beta deviation space, we analyzed the joint distribution of theta and beta z-scores. While a subset of individuals displayed marked increases in theta and reductions in beta power, consistent with the neural slowing profile commonly reported in Parkinson’s disease [71, 72], other patients showed minimal or no deviation from normative centiles. This inter-individual variability was captured in a two-dimensional z-score space, which revealed a continuum of profiles ranging from high-theta/low-beta to low-theta/high-beta. These patterns highlight the substantial heterogeneity in both the direction and magnitude of deviations [71, 73, 74], reinforcing the perspective that Parkinson’s disease is best conceptualized as a spectrum of functional brain alterations rather than a uniform neurophysiological entity.

While this study provides a foundational framework for functional normative modeling using MEG, several methodological limitations merit consideration. From a dataset composition perspective, although our sample is large and drawn from multiple sites, it remains underrepresentative of ethnically and socioeconomically diverse populations, with most participants recruited from Western countries. This limits the global generalizability of the normative reference. Nevertheless, the federated architecture of HBR supports decentralized model updates, enabling the seamless integration of additional datasets without requiring data sharing—a key feature for expanding demographic inclusivity in future work.

In terms of preprocessing, spectral analyses were restricted to the 3–45 Hz range due to limitations of the specparam method [49], precluding investigation of the delta frequency band. Alternative methods for separating periodic and aperiodic activity, such as irregular-resampling auto-spectral analysis (IRASA) [81], may help extend modeling across the full frequency spectrum. Additionally, oscillatory features were computed and averaged at the sensor level to ensure comparability across different MEG systems. While this aggregation reduces spatial specificity, it facilitates device-agnostic harmonization and may enhance transferability to lower-resolution platforms such as EEG or portable devices. Source-space modeling could improve anatomical localization in datasets where co-registered structural MRI is available.

In summary, MEGaNorm represents a major step toward individualized, transdiagnostic analysis of functional brain dynamics across the lifespan. By combining methodological rigor with clinical applicability, our framework lays the foundation for more precise, interpretable, and generalizable models of brain function. As normative modeling continues to evolve, we anticipate that MEGaNorm will serve as a valuable resource for both basic neuroscience and precision neuropsychiatry, helping to bridge the gap between large-scale population data and person-centered clinical insight.

## 4 Methods

### 4.1 Datasets

Resting-state MEG (rs-MEG) data were aggregated from six publicly available datasets, comprising a total of 1,846 clinically undiagnosed participants from the general population (Table 2): the Cambridge Centre for Ageing and Neuroscience (Cam-CAN)[82], Boys Town National Research Hospital (BTH)[36], The Open MEG Archive (OMEGA)[83], the Human Connectome Project (HCP)[84], the National Institute of Mental Health (NIMH)[85], and the Mother Of Unification Studies (MOUS)[86]. Figure 5 shows the age distribution across these datasets. A total of 33 participants were excluded due to missing demographic information, absence of rs-MEG recordings, or convergence failures in the Spectral parameterization algorithm (see Supplementary methods). All included participants were screened for neuropsychiatric conditions and evaluated as typically developing. In addition, we included rs-MEG recordings from 160 patients with Parkinson’s disease available in the OMEGA dataset [83] (67 female; mean age = 59.93 years, SD = 15.59) to demonstrate the clinical applicability of the derived normative models. All patients had a diagnosis of mild to moderate idiopathic Parkinson’s disease and were receiving a stable regimen of antiparkinsonian medication at the time of recording.

**Figure 5:**
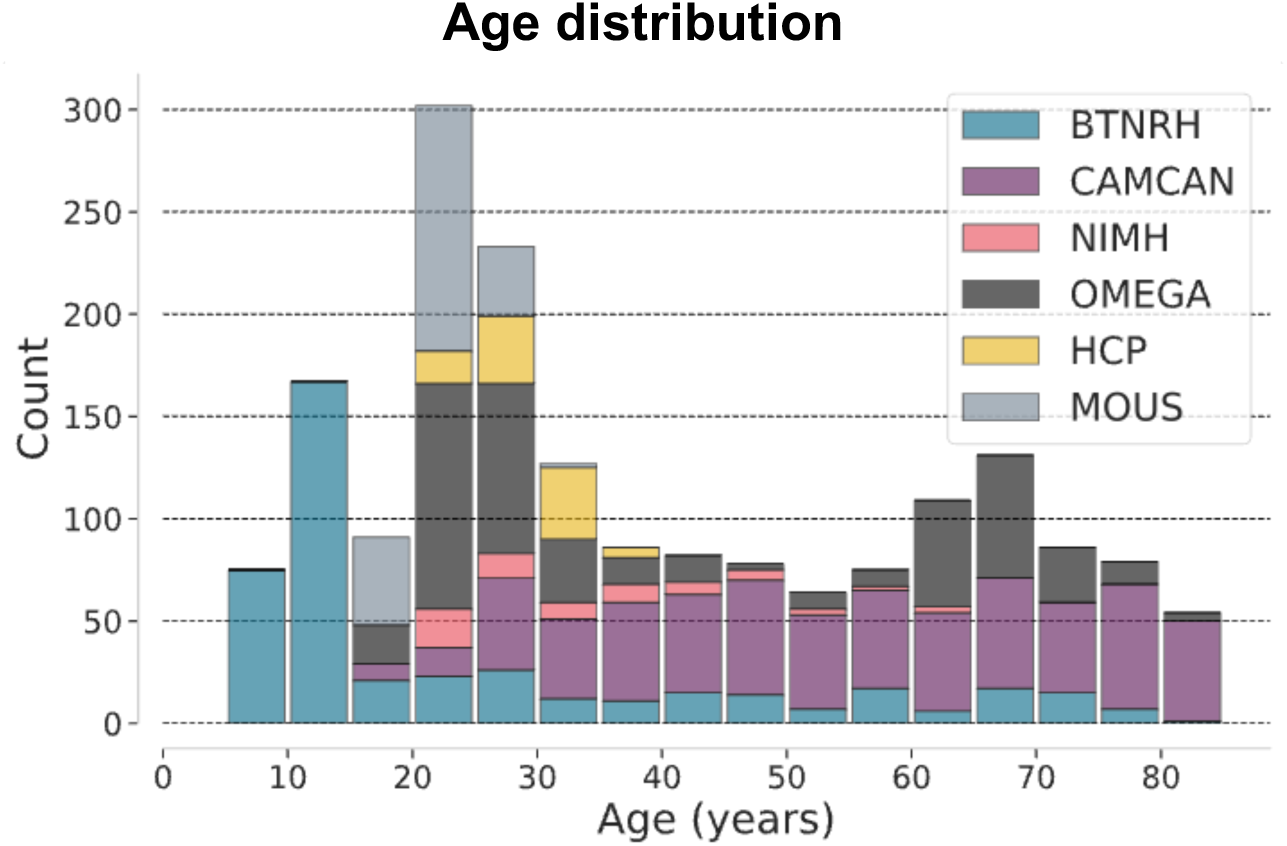
The age distribution of undiagnosed participants across the six datasets. The data spans a broad range of the human lifespan, from childhood (6 years) to late adulthood (88 years).

**Table 2:** Demographics and protocol of the datasets.

| Scanner site | Participants | Female | Mean age | SD | Eyes | Duration | Country |
| --- | --- | --- | --- | --- | --- | --- | --- |
| MOUS [86] | 199 | 101 | 22.02 | 29.47 | EO | 6 min | Netherlands |
| BTH [36] | 433 | 204 | 25.16 | 20.52 | EC | 5–8 min | United States |
| Cam-CAN [82] | 615 | 304 | 54.60 | 18.28 | EC | 8 min 40s | United Kingdom |
| NIMH [85] | 66 | 45 | 34.13 | 12.05 | EC | 6 min | United States |
| OMEGA [83] | 440 | 270 | 41.98 | 20.45 | EO | 5 min | Canada |
| HCP [84] | 89 | 41 | 28.60 | 3.85 | EO | 5 min | United States |

\* Both the number of participants and females are reported after excluding individuals with missing information as described in Methods 4.1; SD = standard deviation; EC = eyes closed; EO = eyes open.

The data acquisition process varied across datasets in terms of MEG hardware configurations, sampling rates, and preprocessing procedures. For the Cam-CAN dataset, recordings were collected using a 306-channel Elekta MEGIN system comprising 102 magnetometers and 204 planar gradiometers, with a 1,000 Hz sampling rate. Real-time head position tracking was performed using four Head-Position Indicators (HPIs), and temporal Signal Space Separation (tSSS) [87] was applied using MaxFilter v2.2 (correlation threshold = 0.98, 10-second sliding window) to mitigate motion artifacts and external noise. Electrocardiogram (ECG) and electrooculogram (EOG) signals were also recorded to capture physiological activity. The BTH dataset was acquired using a similar Elekta MEGIN system (1,000 Hz sampling rate) with four HPIs, and tSSS was applied using MaxFilter v2.2 (correlation threshold = 0.95, 10-second sliding window). The OMEGA dataset utilized a 275-channel CTF axial-gradiometer system (CTF MEG,

Coquitlam, BC, Canada) with a 2,400 Hz sampling rate, along with ECG and EOG recordings, and applied third-order gradient noise correction during acquisition to reduce environmental interference. The MOUS dataset was also acquired using a CTF system, with a 1,200 Hz sampling rate and accompanying ECG and EOG channels. Similarly, the NIMH dataset employed a CTF system with a 1,000 Hz sampling rate and third-order gradient noise correction. Finally, the HCP dataset was recorded using a Magnes 3600 system (4DNeuroimaging, San Diego, CA) with 248 magnetometers.

### 4.2 MEG preprocessing

MEG recordings were preprocessed using a pipeline designed to reduce noise and prepare the data for spectral analysis while accommodating variability in acquisition protocols across sites. Preprocessing began with the removal of bad channels to ensure signal quality before further analysis. First, channels previously marked as flat or noisy in the original datasets were excluded. For the Cam-CAN and BTH datasets, which were recorded using Elekta MEGIN systems, we used the MaxFilter-processed data after application of tSSS. In these datasets, noisy channels had already been identified and either reconstructed or excluded during acquisition. For the remaining datasets, recorded using CTF or Magnes systems, bad channel detection was performed using Maxwell filtering as implemented in MNE-Python [88]. Channels identified as excessively noisy or flat were removed before subsequent steps.

All recordings were resampled to 1,000 Hz if necessary and filtered using a finite impulse response (FIR) filter with a 1 Hz high-pass and 45 Hz low-pass cutoff. A notch filter was applied to attenuate line noise. To reduce physiological artifacts, independent component analysis (ICA) was performed using the FastICA algorithm [89], as implemented in MNE-Python [88], retaining 30 principal components. When ECG recordings were available, Pearson correlation coefficients were calculated between each independent component and the ECG signal to identify components related to cardiac activity. In the absence of ECG data, a synthetic ECG reference was generated by averaging across magnetometers (for Elekta and Magnes systems) or gradiometers (for CTF systems). Components with correlation coefficients exceeding 0.9 were rejected. A similar procedure was used to identify and remove eye movement artifacts when EOG recordings were available. To minimize contamination from non-neural transients such as eye blinks or movements typically occurring at the start and end of sessions, the first and last 20 seconds of each recording were discarded. The remaining data were segmented into 10-second epochs with a 2-second overlap, creating standardized time windows for downstream spectral analysis.

### 4.3 Estimating power spectrum densities and f-IDP extraction

To extract functional imaging-derived phenotypes (f-IDPs), we first computed power spectral densities (PSDs) for each 10-second segment using Welch’s method [90], employing a 2-second Hamming window with 1-second overlap and no zero-padding. The resulting PSDs were then averaged across all segments to yield a single PSD estimate per sensor, thereby enhancing the signal-to-noise ratio.

MEG recordings reflect both oscillatory (periodic) brain activity and broadband aperiodic signals. Consequently, raw PSDs can be confounded by background 1*/f* like structure (Figure 6a), where spectral power decays exponentially with frequency *f* [49]. To prevent the confounding effect of aperiodic activity, we first isolated periodic activity by subtracting the aperiodic activity (Figure 6b). This ensures that the extracted f-IDPs reflect true oscillatory brain activity. We then computed the relative periodic power for canonical frequency bands: theta (3–8 Hz), alpha (8–13 Hz), beta (13–30 Hz), and gamma (30–40 Hz) (Figure 6c). Relative power in each band was defined as the ratio of periodic power within that band to the total periodic power across the full 3–40 Hz range.

**Figure 6:**
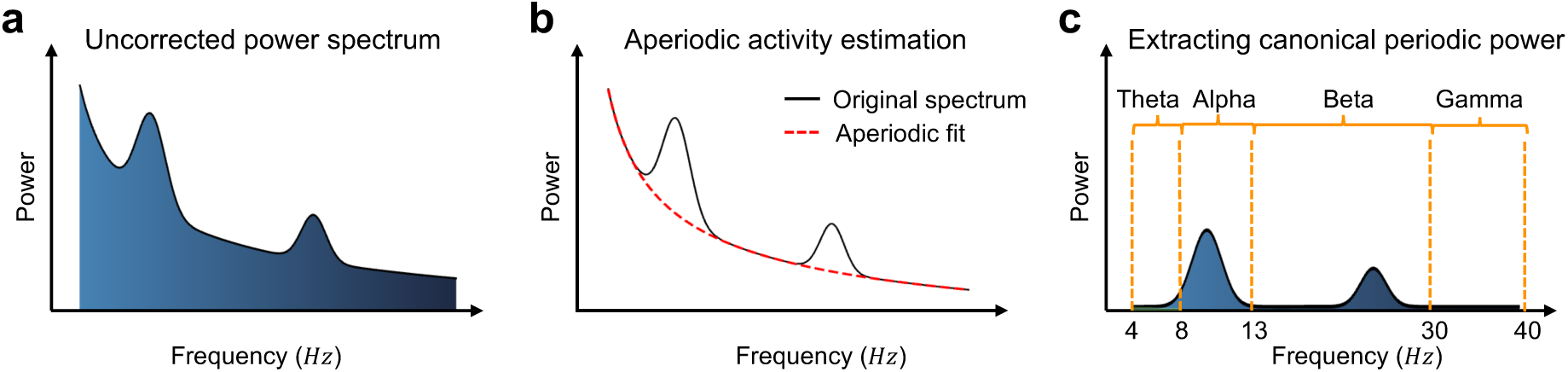
Extracting relative power of four canonical frequency bands. **(a)** The original power spectral density (PSD), encompassing both aperiodic and periodic activities. Spectral peaks reflect brain oscillations, while the broadband 1*/f* activity corresponds to the aperiodic component. **(b)** The aperiodic component (red dashed line) is estimated using an exponential function from the original PSD within the Spectral parameterization (specparam) algorithm. The aperiodic activity is then subtracted from the original PSD to isolate periodic activity. **(c)** Relative power of each canonical frequency band (theta, alpha, beta, gamma) is computed from the periodic activity as a proportion of the total periodic power between 3–40 Hz.

To isolate the periodic component, we used the Spectral parameterization (specparam, formerly fooof) algorithm (version 1.1.0)[38], which decomposes the PSD into periodic and aperiodic components (Figure 7b) by modeling the latter as an exponential function:

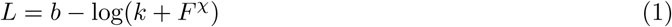

where *b*, *F*, and *χ* represent the broadband offset, frequency vector, and exponent, respectively. The parameter *k* captures the bend (or ”knee”) in the aperiodic component.

**Figure 7:**
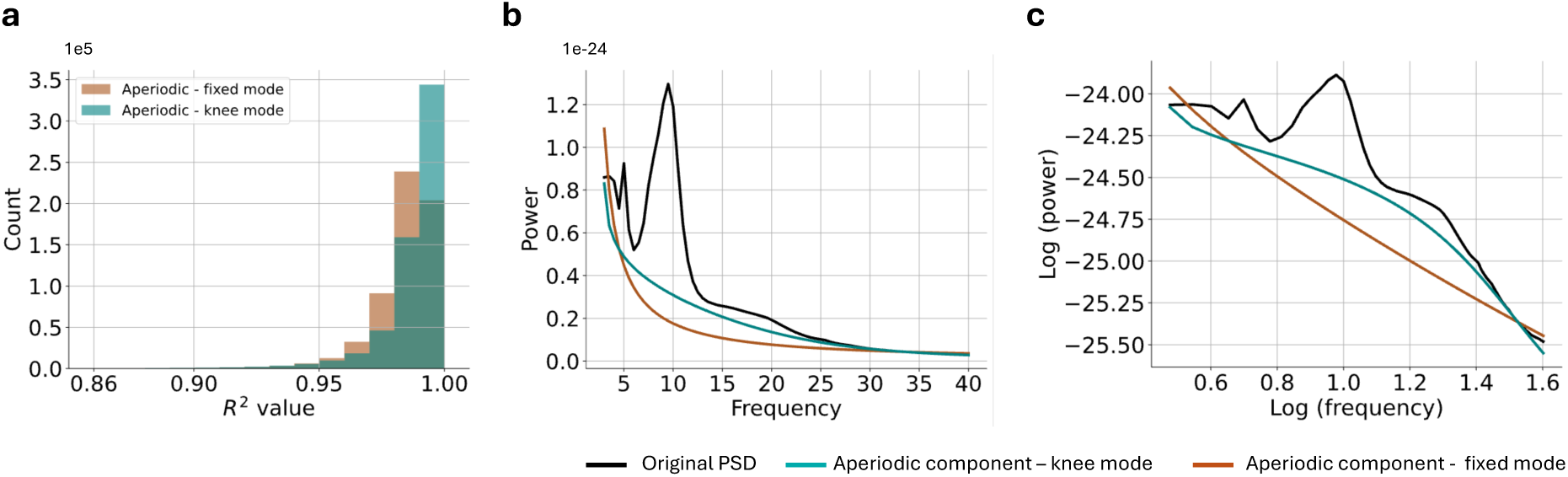
Parametrizing the bend (”knee”) in aperiodic spectra improves specparam model fits. **(a)** Histogram of the coefficient of determination (*R*^2^) for model fits across participants in both fixed mode (*k* = 0) and knee mode (*k >* 0). The knee mode yields a slightly higher *R*^2^ value (median=0.99, SD=0.018) compared to the fixed mode (median=0.98, SD=0.019). **(b)** Average power spectrum across channels and participants (black line), alongside the corresponding estimated aperiodic components using both knee (teal line) and fixed (brown line) parameters, shown in linear space. At lower frequency ranges, the fixed-mode aperiodic fit overshoots the original PSD. **(c)** The same data shown in log-log space. The original PSD displays a clear bend in the middle frequency ranges. The fixed-mode fit (brown) fails to capture this bend, appearing as a straight line, while the knee-mode fit (teal) successfully models the curvature.

Incorporating the knee parameter (*k >* 0) into the fitting process significantly enhances the quality of specparam model fits compared to the fixed mode (*k* = 0). Figure 7a shows that enabling the knee mode substantially increases the proportion of explained variance (*R*^2^). When the model is fit in fixed mode, the aperiodic component can exceed the original power spectrum in certain frequency ranges, resulting in biologically implausible negative periodic power values (Figure 7b). Moreover, the average power spectrum across subjects and channels, plotted in log-log space, reveals a clear knee in the mid-frequency range (Figure 7c). The fixed mode, which imposes a single 1*/f* characteristic, appears as a straight line in log-log space and fails to capture this knee. In contrast, enabling the knee mode parameter allows models to more accurately represent this spectral bend, improving the quality of the fit.

To ensure the goodness of fits, the coefficient of determination *R*^2^ is used as a quality control approach. This metric quantifies how much of the shape of the spectrum is captured by specparam’s model. We excluded channels with *R*^2^ *<* 0.90 from further analysis to ensure reliable aperiodic correction.

To obtain the f-IDPs, we spatially averaged them across all MEG sensors. This summarization step yielded four global f-IDPs per participant, each representing the relative periodic power in one of the canonical frequency bands across the entire brain. For recordings from Elekta MEGIN systems, we computed the average across both magnetometer and planar gradiometer channels, as these sensor types showed strong agreement in their spectral profiles (Pearson correlation; see Supplementary results). This sensor-level aggregation was necessary for harmonizing across recording devices and enabling consistent downstream modeling of spectral features. We use the MEGaNorm package (v0.1.0)[65] for MEG data processing and f-IDP extraction.

### 4.4 Normative modeling

We used hierarchical Bayesian regression (HBR), implemented in the Predictive Clinical Neuroscience Toolkit (PCNtoolkit, version 0.35)[91], to estimate normative ranges for the extracted f-IDPs. HBR was selected for its ability to account for site-related batch effects and its compatibility with federated learning architectures[4], enabling decentralized model updates without sharing individual-level data. Age was included as a continuous covariate, and to model non-linear age trajectories of f-IDPs, we applied a cubic B-spline basis expansion with five evenly spaced knots. Sex and site were modeled as random intercepts to account for systematic differences in baseline power levels across subgroups. To further accommodate heteroscedasticity, the variance was modeled as a function of age, allowing for age-dependent variability in the distribution of f-IDPs.

To accommodate non-Gaussian distributions in the data, we used a Sinh-Arcsinh (SHASH) likelihood [39], which generalizes the Gaussian distribution by introducing additional parameters for skewness and kurtosis. The SHASH distribution [42] is parameterized by four quantities: the mean (*µ*), scale (*σ*), skewness (*ɛ*), and kurtosis (*δ*), allowing flexible modeling of both the location and shape of the distribution (see Supplementary methods). In our implementation, all four parameters were modeled as linear functions of age, enabling conditional estimation of both central tendency and distributional form. Posterior inference was performed using the No-U-Turn Sampler (NUTS)[92], as implemented in PyMC [93] version 5.22.

### 4.5 Model evaluation

To assess the goodness of fit and calibration of the estimated normative centiles, we evaluated the trained models on a held-out test set using a range of diagnostic metrics. The dataset was split into 50% training and 50% testing subsets stratified by sites. This procedure was repeated 10 times using different random seeds to assess the robustness and generalizability of the models.

We evaluated both the accuracy of the fit and the quality of centile calibration, following established principles in normative modeling [94]. For model fit, we used the Standardized Mean Squared Error (SMSE) to quantify prediction accuracy. To assess the distributional correctness of the estimated centiles, we examined the Gaussianity of the derived z-scores using three complementary metrics: skewness, excess kurtosis, and the Shapiro–Wilk test statistic (*W*) [43]. These metrics provide an indirect measure of centile quality by testing whether the z-scores conform to a standard normal distribution.

To directly evaluate centile calibration, we introduced a new metric, Mean Absolute Centile Error (MACE), which quantifies the average deviation between predicted and empirical centiles in the test set. Inspired by reliability diagrams, MACE measures the alignment of model-predicted quantiles with observed data quantiles (see Figure 8). The MACE metric is computed as follows:

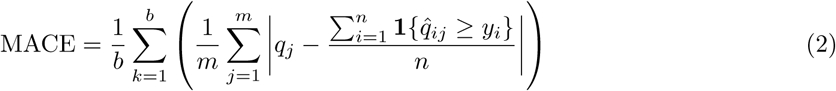

where *b* represents the number of batch effects (i.e., unique combinations of sex and site), and *m* denotes the number of centiles used for calibration. The terms *q_j_* and *q*^*_ij_* represent the centile value and its corresponding predicted y-value for the *i*th data point, respectively. The term **1***{q*^*_ij_ ≥ y_i_}* is an indicator function that outputs 0 or 1, depending on whether *y_i_* lies below or above its predicted *j*th centile value. Summing these outputs across all data points and dividing them by the total number of data points, *n*, gives the actual centile corresponding to the *j*th predicted centile. By subtracting this from the true *j*-th centile, we can have an estimation of the deviation between the predicted and true centiles. In our experiments, we evaluate the MACE for the 1^st^, 5^th^, 25^th^, 50^th^, 75^th^, 95^th^, and 99^th^ centiles.

**Figure 8:**
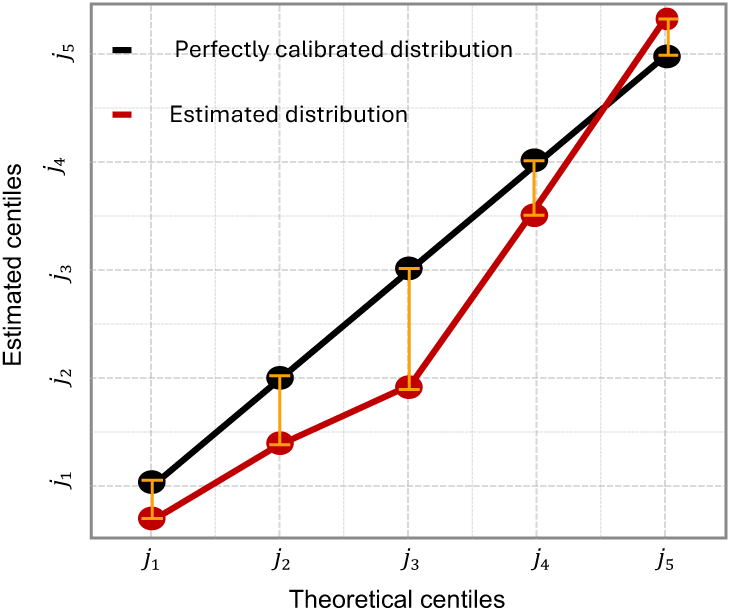
Schematic illustration of calculating the Mean Absolute Centile Error (MACE) to evaluate the quality of estimated centiles. The Reliability plot. The x-axis represents the *i*-th theoretical centile, while the y-axis shows the corresponding estimated centile. A well-calibrated model should align with the 1:1 diagonal line (shown in black). In this example, the estimated centiles (shown in red) deviate from the expected line. In MACE, we compute the difference between the theoretical and actual centiles for each *j*-th centile (orange lines) and then average these differences across all centiles.

### 4.6 Anomaly detection

We assessed the clinical relevance of the derived normative models using an anomaly detection framework, in which large deviations from normative trajectories are interpreted as potential markers of pathology. Deviation scores (*z*) were computed for all patients and healthy participants in the test set of the OMEGA dataset. Following the approach proposed by Kia et al. [4], each *z*-score was converted into a probability of abnormality, *P*_abn_(*z*), by calculating the area under the standard Gaussian cumulative density function in the interval [*−|z|, |z|*]:

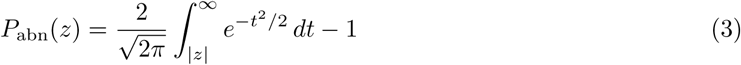

This transformation yields a continuous measure of abnormality for each f-IDP. To quantify discriminative performance, we computed the area under the receiver operating characteristic curve (AUC) for each feature, benchmarking abnormality probabilities against true clinical labels across test samples.

## Code Availability

All custom code developed for data processing, model training, and analysis is openly available in the MEGaNorm repository at https://github.com/ML4PNP/MEGaNorm and archived on Zenodo [65]. The code is released under the GNU General Public License v3.0 and includes documentation for installation and usage. Scripts to reproduce the main analyses and figures in this paper are provided in a separate repository at https://github.com/ML4PNP/MEG_Norm. Additionally, we plan to openly share the derived normative models via the PCNPortal at https://pcnportal.dccn.nl/ for model extension and adaptation to new datasets.

## Data Availability

All datasets used in this study are publicly available from established open-access neuroimaging repositories. Below is a summary of the data sources and access links: The HCP dataset [84] is available from the Human Connectome Project repository: https://www.humanconnectome.org/study/hcp-young-adult. The Open MEG Archive (OMEGA) dataset [83] can be accessed via https://doi.org/10.23686/0015896. The National Institute of Mental Health (NIMH) dataset [85] is hosted on OpenNeuro at https://doi.org/10.18112/openneuro.ds004215.v1.0.024. The Cambridge Centre for Ageing and Neuroscience (Cam-CAN) dataset [82] is available through the CamCAN data portal: https://camcan-archive.mrc-cbu.cam.ac.uk/dataaccess/. The Boys Town National Research Hospital (BTH) dataset [36] can be downloaded via the link provided in the original publication: https://cdn.boystown.org/media/Rempe_Ott_PNAS_2023_Data.zip. The Mother Of Unification Studies (MOUS) dataset [86] is available from the Radboud Data Repository: https://doi.org/10.34973/37n0-yc51. We gratefully acknowledge the considerable open-science efforts of the neuroimaging community in making these datasets publicly available. This work would not have been possible without the commitment of these research teams to data sharing and transparent science.

## Competing interests

The authors declare no competing interests.

## Acknowledgments

S.M.K. gratefully acknowledges the starter grant for the *MEGaNorm* project, funded by the Dutch Ministry of Education, Culture and Science under the National Sector Plan. S.M.K. further acknowledges support from the Netherlands Organization for Scientific Research (NWO) through the Small Compute Applications grant (EINF-8659) for the project *Charting Functional Brain Activities*. S.M.K. thanks the Digital Sciences for Society program at Tilburg University for the Growth Project grant (DSFS 202417) supporting the project *Charting the Normative Electroencephalography in Healthy Aging Population*.

## Notes

https://github.com/ML4PNP/MEGaNorm

### Competing Interest Statement

The authors have declared no competing interest.

