## Supplementary Materials for "MEGaNorm: Normative Modeling of MEG Brain Oscillations Across the Human Lifespan"

#### Supplementary Methods

##### The SHASH likelihood

The SHASH distribution [1] incorporates four key parameters—mean ($\mu$), variance ($\sigma$), skewness ($\epsilon$), and kurtosis ($\delta$), when defining the likelihood in the Bayesian modeling framework [2]. Unlike the Gaussian distribution ($X \sim N(\mu, \sigma)$), which depends solely on the location $\mu$ and scale $\sigma$ parameters, the SHASH distribution models also the shape of the distribution by applying an inverse sinh-arcsinh transformation to samples drawn from a standard Gaussian distribution ($Z \sim N(0, 1)$):

$$\xi^{-1}_{\epsilon,\delta}(z) = \sinh\left(\frac{\sinh^{-1}(z) + \epsilon}{\delta}\right)$$

Here, $z$ represents samples from a standard Gaussian distribution, and the transformation results in a SHASH distribution $S(\epsilon, \delta)$, where the parameters $\epsilon$ and $\delta$ control the skewness and kurtosis of the distribution, respectively. Specifically, $\epsilon$ adjusts the asymmetry of the distribution (skewness), while $\delta$ governs the tail behavior (kurtosis), allowing the SHASH distribution to model a wide range of non-Gaussian data shapes. The location ($\mu$) and scale ($\sigma$of the distribution are also incorporated as follows:

$$\Omega = \xi_{\epsilon,\delta}^{-1}(Z)\sigma + \mu$$

where $\Omega \sim \mathcal{S}(\mu, \sigma, \epsilon, \delta)$. Together, the four parameters enable the SHASH distribution to accurately capture complex characteristics of data distribution, such as non-Gaussianity, and varying skewness or kurtosis.

#### Excluded participants

Some participants were excluded from the analysis for the following reasons: missing demographic information, missing magnetoencephalography (MEG) recordings, and failure to fit models using the spectral parameterization algorithm. The table below summarizes the number of excluded participants per dataset:

**Supplementary Table** 1: Number of excluded participants per dataset.

| Scanner site | Number of excluded participants |
|---|---|
| BTH | 0 |
| CamCAN | 20 |
| NIMH | 0 |
| OMEGA | 2 |
| HCP | 6 |
| MOUS | 5 |

### Supplementary Results

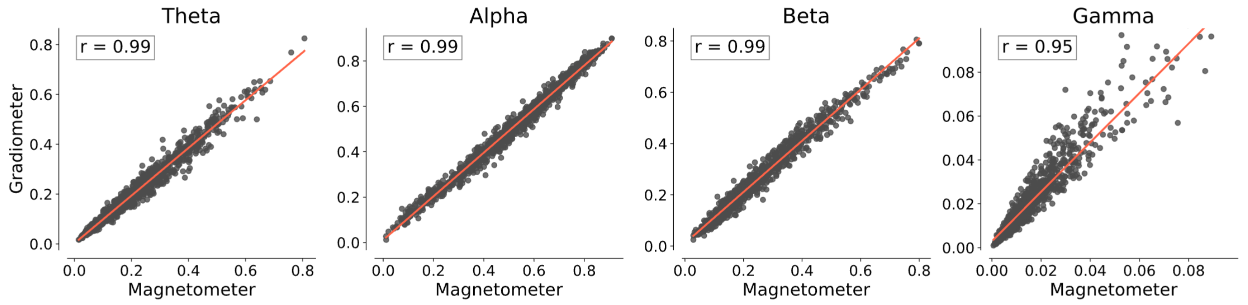


**Supplementary Figure** 1: The relative power of canonical frequency bands exhibited a high Pearson correlation across magnetometers and gradiometers. All Pearson correlation coefficients ($r$) were greater than 0.95.

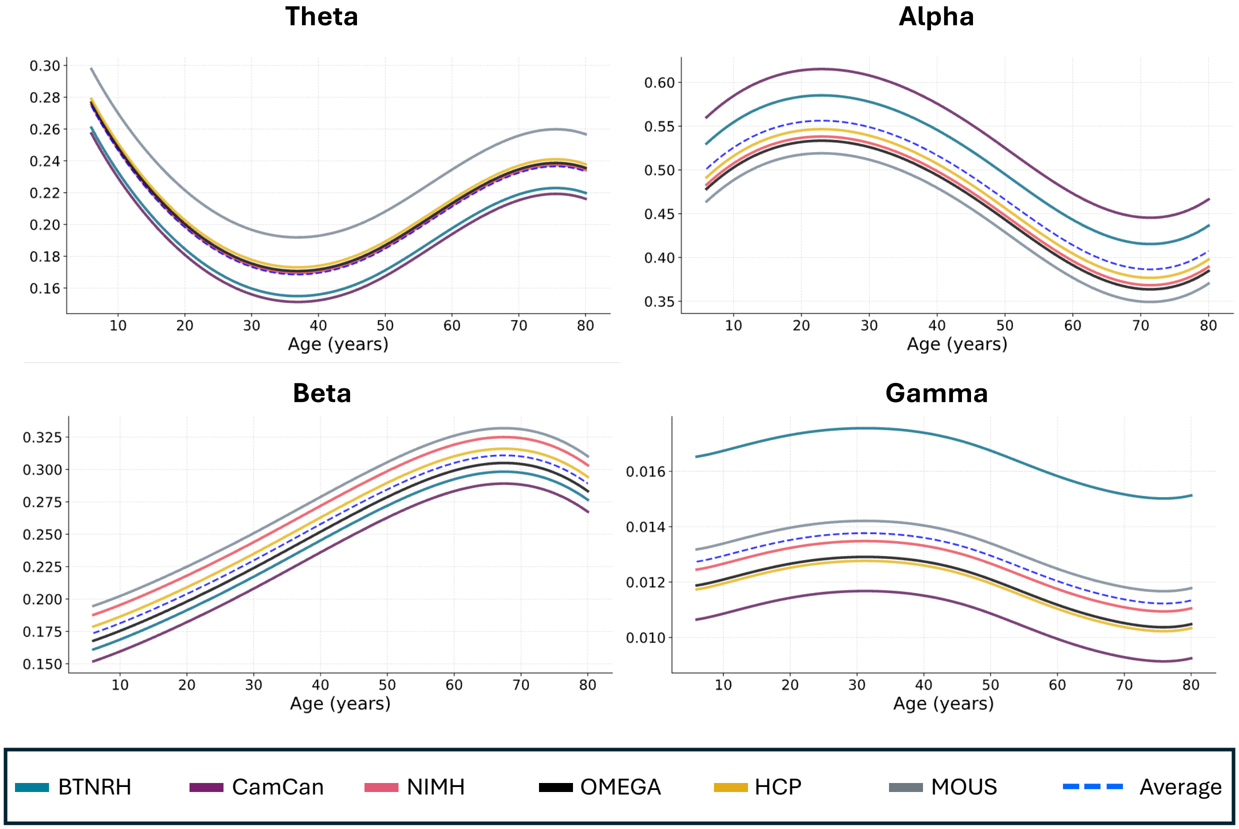


**Supplementary Figure** 2: Lifespan $50^{th}$ centile trajectory of f-IDPs in males across six sites. The $50^{th}$ centile for each site is estimated by drawing samples from the posterior predictive distribution. Additionally, the average trajectory across sites is represented by a blue dashed line. The MOUS, OMEGA, and HCP datasets that were recorded in the eyes-closed condition exhibit a lower relative alpha power compared to the average.

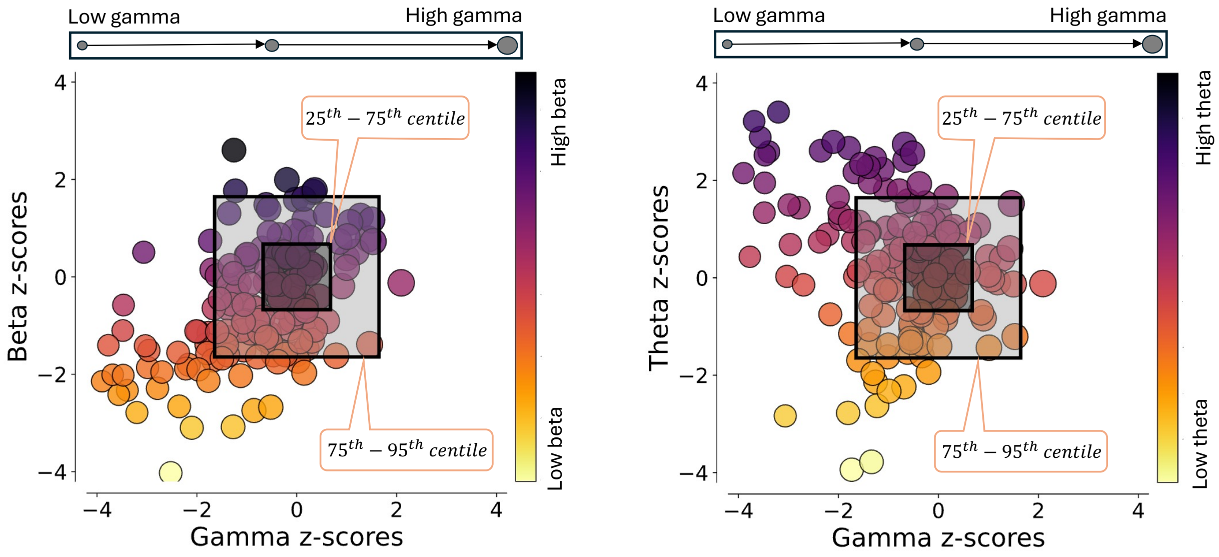


**Supplementary Figure** 3: Scatter plot of the distribution of Parkinson's disease patients in gamma-beta and gamma-theta deviation space. The scatter plots highlight the heterogeneity in the patient population as spectra. Marker size represents the x-axis values, while the color map corresponds to the y-axis values.
